# Over the hills and far away: linking landscape factors with cavity excavation on living forest trees by the Black Woodpecker (*Dryocopus martius*, L. 1758)

**DOI:** 10.1101/2022.06.22.497197

**Authors:** Cedric Cabrera, Jean-Matthieu Monnet, Jean-Jacques Boutteaux, Baptiste Doutau, Pascal Denis, Yoan Paillet

## Abstract

The Black Woodpecker (*Dryocopus martius*, L. 1758) is the largest primary cavity excavator in Europe. Its cavities represent an essential microhabitat for many other forest species and the knowledge on landscape factors linked with cavity excavation by the Black Woodpecker is needed to support the conservation of this species and associated species. Such relationships should thus be quantified at different scales ranging from the stand to the extended home range.

We used cavity maps established by foresters and naturalists to build a large (2689 cavity bearing trees) database distributed over several sites in France. Based on this and on a set of background points, i.e. randomly selected points devoid of cavity in the vicinity, we analysed the effects of stand composition and landscape features (forest cover, forest connectivity and fragmentation) at three different scales around each cavity and background point corresponding to a forest management unit (10ha), the core (100ha) and extended (250ha) home range scales.

We showed that indices describing forest continuity (cohesion, landscape shape index) and forest tree species composition (especially the presence of mixed forests) had significant positive effects but that the magnitude varied across the three scales. We notably observed the strongest effects at the core home range scale (100ha), indicating that Black Woodpecker requirements for cavity excavation are more pronounced at this scale. The Black Woodpecker tends to avoid pure conifer-dominated stands to excavate cavities, but benefits from mixed forests, that couple favourable foraging and cavity excavation sites. The bird also prefers continuous forest landscapes with high cohesion and low edge densities. We also showed that the positive effects of forest landscape were generally at higher elevation, indicating context-dependence.

Forest planning rarely integrates the landscape patterns. A better understanding of the features linked with cavity excavation by the Black Woodpecker may hence help to better integrate their conservation in forest management planning. Our results also show the importance to maintain mixed broadleaf-conifer forests as well as continuous and well-connected forest landscapes to favour features that benefit primary and secondary cavity nesters.

## Introduction

The conservation of forest biodiversity relies on an ensemble of methods including setting aside forest reserves and small conservation patches and preserving, within managed forests, elements that favor the biodiversity of forest dwelling species (Bollmann and Braunisch 2013; Kraus and Krumm 2013). Tree-related microhabitats (hereafter microhabitats, sensu Larrieu et al. 2018), defined as tree-born singularities, are structural elements that support various species and communities. Among the different microhabitats, woodpecker cavities are, so far, the most documented regarding the biodiversity they harbor (Cockle et al. 2011; Larrieu et al. 2018). Black Woodpecker (*Dryocopus martius* L. 1758) cavities are the largest excavated by a bird in Europe, and, as such, they have a crucial role for many secondary cavity nesters (e.g. Johnsson 1993). The Black Woodpecker excavates for nesting, but also for roosting, and both cavity types may be used by other nesters. The Black Woodpecker is qualified as an ecosystem engineer, because it significantly modifies its environment and thus allows other species to live and reproduce (Johnsson 1993; Kosiński et al. 2010; Pirovano and Zecca 2014): e.g. the Boreal Owl (*Aegolius funereus*, L. 1758), the Stock Dove (*Columba oenas*, L. 1758), the Greater Noctule bat (*Nyctalus lasiopterus*, S. 1780), the Pine Marten (*Martes martes*, L. 1758), and numerous saproxylic beetles and fungi. This effect is accentuated by the fact that this bird has a large home range (316ha in the Italian Alps, Bocca et al. 2007; Rolstad et al. 1998; up to 500ha in Sweden and Norway, Tjernberg et al. 1993), and it may contribute to increased diversity of niches over large areas. It also needs forests with old and large trees to accomplish its life cycle (Basile et al. 2020; Khanaposhtani et al. 2012; Olano 2015). Understanding the factors linked with cavity excavation by the Black Woodpecker is essential to better integrate them in forest management at different scales. However, such knowledge relies on expertise (e.g. Cuisin 1967) and has seldom been quantified for western Europe using systematic approaches on large datasets (see Brambilla and Saporetti 2014; Gil-Tena et al. 2013; Tjernberg et al. 1993).

At the tree scale, several studies have shown that the Black Woodpecker tend to excavate in less crowded environments (i.e. with less undergrowth), on trees devoid of low branches and trees with a diameter at breast height larger than the others in the vicinity (Basile et al. 2020; Puverel et al. 2019). Some papers also mention fungi decay as a strong facilitator of woodpecker excavation (Jackson and Jackson 2004; Zahner et al. 2012). At larger scales, several stand and landscape factors are also linked with cavity excavation, notably forest fragmentation and landscape heterogeneity (Bełcik et al. 2020; Brambilla and Saporetti 2014; Gil-Tena et al. 2013; Saporetti et al. 2016). The Black Woodpecker depends on forest habitats to reproduce (Angelstam et al. 2002; Mikusinski 1995; Tjernberg et al. 1993), but also on more open forest habitats like clearings to feed (Brambilla and Saporetti 2014; Dorresteijn et al. 2013; Pirovano and Zecca 2014). Forest connectivity would positively influence the Black Woodpecker presence and cavity excavation (see e.g. Karimi et al. 2018 for an example in Iran), with higher probabilities in places where favorable patches are connected (Gil-Tena et al. 2013; Tobalske and Tobalske 1999). Although the Black Woodpecker tends to prefer broadleaves to nest in Western Europe (Mikusiński 2015), it feeds on insects – ants and beetles larvae – found mostly in coniferous forests (Mikusiński 1997). On the top of that, the Black Woodpecker shows behavioral variations – context dependence – across its distribution area, notably in terms of nesting and excavating preferences (Mikusiński 2015), suggesting idiosyncrasies linked to the climate (precipitation, temperature) and topography (elevation).

In this context, we aimed to quantify the effects of forest tree species composition and landscape configuration on cavity excavation by the Black Woodpecker, and ultimately to better integrate cavities conservation in forest management and planning. We gathered a large database of Black Woodpecker cavities maps provided by local managers and naturalists on different sites in France covering a wide range of ecological conditions (forest types, elevation, climate). We worked at three different scales to provide management and biologically relevant answers: (i) 10ha (0.1km2) scale that corresponds to a typical forest management unit in France; (ii) 100ha (1km2) scale that corresponds to the mean core home range of the Black Woodpecker where it defends its territory and breeds (Bocca et al. 2007); (iii) 250ha (2.5km2) scale that corresponds to the mean home range over the year (Olano 2015). For each scale, we extracted the same landscape variables to assess which ones had most influence on cavity excavation, and understand whether the landscape factors were the same across all scales. To control for potential effect of the context on these responses, we also included biogeographical and climatic variables to the analyses (elevation, aspect, temperature, precipitations). We hypothesized that:

i. The presence of both broadleaf stands for nesting and conifer stands for feeding would positively influence the presence of cavities;
ii. The quantity of large, continuous, forest areas would positively influence the presence of cavities. This would consist in zones with high forest cover and high values of connectivity between patches;
iii. The landscape heterogeneity, measured as the size and distribution of forest and non-forest patches, would negatively influence the presence of cavities;
iv. These relationships vary according to the context, notably between lowland and mountain forests.

## Materials and methods

### Study sites and dataset

We worked in five French forest sites where Black Woodpecker cavities have been mapped by forest managers and naturalists (Figure 1).

**Figure 1:**
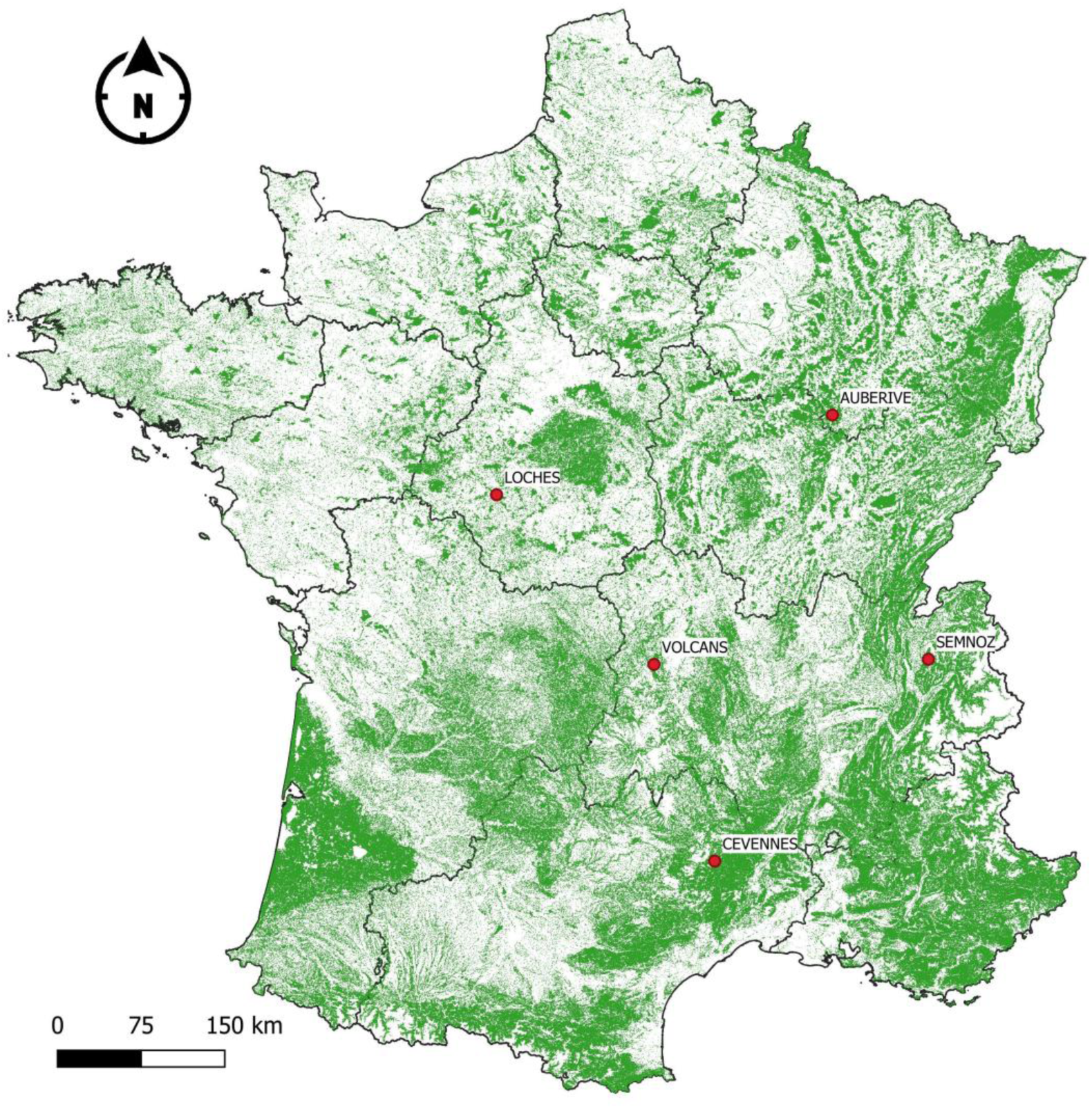
Map of the study sites. Green layer represents forest cover.

The dataset has been compiled from different sources (National Forest Service, non-governmental organizations, local naturalists, National and Regional Parks) but originates from comparable field campaigns and methods: Black Woodpecker cavities were systematically and actively searched on all sites either during dedicated field campaigns or other forestry activities (e.g. tree-marking operations, see Table 1 for specifications on methods and data providers). Mapping cannot be assumed to be fully exhaustive and some cavities may have been missed by the inventories, but regarding the quantity of cavities mapped, and the methods used to analyze these data (see below), we remain confident that our results highlight unbiased trends on the link between cavities and landscape features. Most cavities were observed from the ground, without relying on pole-cameras or other devices except binoculars (see discussion for the potential bias linked to this method of observation). The inventories were reliable since Black Woodpecker cavities cannot be confounded in the field with other woodpecker cavities (Larrieu et al. 2018) and professional birders and naturalists were involved in the mapping process. It is important to note that we did not consider only breeding cavities, but all cavities excavated by the bird. The presence of such cavities (whatever their use by the Black Woodpecker) in the landscape is crucial for many species and studying landscape drivers of their excavation is valuable in terms of management. In addition, we checked a small sample of the cavities in the field using pole cameras for a side project. We searched about 80 cavities distributed over the 5 sites (in 2022 and 2023), and it appeared that around 13% could not be confirmed because of tree harvesting or breakage. All other trees could be found in the field and corresponded to actual cavities.

**Table 1:** Distribution and characteristics of Black Woodpecker cavity locations selected for the study.

| Site | Forest type | Cavity number | Elevation (m) |  | Mean annual temperature (°C) |  | Mean annual precipitation (mm) |  | Slope (deg) |  | Methods of cavity inventory | Data provider |
| --- | --- | --- | --- | --- | --- | --- | --- | --- | --- | --- | --- | --- |
|  |  |  | Min/Max | Mean | Min/Max | Mean | Min/Max | Mean | Min/Max | Mean |  |  |
| Auberive | Mixed beech-oak | 194 | 360/504 | 440 | 8.7/9.6 | 9.1 | 802/911 | 853 | 0/31.2 | 7.8 | Inventory during tree selection operations | National Forest Service (ONF) |
| Cévennes | Mixed beech-fir | 2094 | 796/1400 | 1151 | 5.9/10.2 | 7.6 | 774/979 | 868 | 1,1/48.1 | 18.35 | Active and systematic search by foresters and National Park staff | National Forest Service (ONF) and Cévennes National Park |
| Loches | Oak dominated | 111 | 91/141 | 124 | 11/11.3 | 11.1 | 690/716 | 703.8 | 0/9.7 | 1.9 | Active searching by a naturalist | M. Dubois (naturalist) |
| Semnoz | Mixed beech fir | 37 | 1017/1500 | 1353 | 4.6/7.5 | 5.6 | 1154/1430 | 1335 | 4.3/28 | 12.2 | Active and systematic searching by a NGO | Ligue pour la Protection des Oiseaux (LPO, B. Doutau) |
| Volcans | Mixed beech fir | 252 | 685/1154 | 935 | 6.5/8.7 | 7.6 | 765/912 | 841 | 0.9/40.7 | 10.3 | Active and systematic searching by a regional park | Parc Naturel Régional des Volcans d'Auvergne |

We harmonized the different datasets and selected data to control for dependence and heterogeneity. We first excluded observations made before 2010 (dates of all observations 2000 – 2019), since we considered that landscape (forest) variables could have changed significantly since then, because of e.g. tree mortality and anthropogenic or natural disturbances (e.g. harvesting, tree dieback, storms).

Cavities were rarely observed on dead trees and snags since most managers target living trees for forest management purposes (in total 18 cavities were reported on dead trees on the initial dataset). Although they constitute quality breeding and foraging sites for the birds, dead trees and snags are relatively ephemeral habitats that can be more easily missed out by the inventories. We excluded the few cavities on dead trees since the conditions of excavation may rely more on tree vitality than landscape in this case and it is also difficult to assess whether a cavity on a dead tree was excavated before or after the tree died (Puverel et al. 2019). In addition, it is likely that when the Black Woodpecker has the choice of excavating site, especially when smooth-bark broadleaves are present, the dead trees are not privileged over living trees, so that in Western Europe the prevalence of cavities on dead trees is lesser than in other, conifer-dominated, contexts such as Eastern Europe or Fennoscandia (Zawadzki 2023; Zawadzki and Sławski 2023). For these reasons, we assume that our sample is not systematically biased by living trees.

Based on this first selection, we kept 2689 Black Woodpecker cavities observations with spatial coordinates (Table 1). As indicated by the data providers, all locations were measured with commercial-grade GPS receivers or directly plotted on maps. Precision is thus comprised between 10 m and 100 m, which is consistent with the minimum scale of analysis (10ha).

The dataset is presence-only data, so we generated background points to process statistical analyses (Renner et al. 2015). Random generated background points is a powerful method to deal with presence-only data, notably because background points are less time-consuming than true absence data to gather, even if they are sometimes less precise (e.g. Iturbide et al. 2015). We preferred this method over other presence only analysis methods (see background points selection below) because it allowed a better control of the location and distribution of background points.

We selected 5km radius areas around the presence locations and kept only forested areas (from the satellite images derived layer OCSOL 2018, https://www.theia-land.fr/ceslist/ces-occupation-des-sols/, Inglada et al. 2017). Then, we excluded an area of 250m radius around each presence location to limit the spatial auto-correlation between presence and background point locations. Although this distance does not insure independence of the observations (250m radius corresponds to approximately 20ha), it represents a compromise between spatial autocorrelation and power of the analyses.

We then compiled 2689 presence data with approximately ten times more (22326) background points drawn at random in the envelopes described above, for a total of 25015 locations. To ensure spatial independence between the observations, we used spatial thinning to conserve presence data that were distant of at least 250m (i.e. we randomly selected locations that were at least 250m apart), and selected ten-times more background point locations within each site (R package spThin, Aiello-Lammens et al. 2015). The training dataset finally comprises 7348 locations (668 presence and 6680 background points).

We then delineated the three buffer zones corresponding to our three study scales (radius 179m (10ha); radius 565m (100ha) and radius 879m (250ha) around each location) and extracted forest, landscape and biogeoclimatic variables (Table 2).

**Table 2:** Explanatory variables and sources used to characterize the environment of Black Woodpecker cavities and background points. Variables definitions and sources are detailed in the text.

| Group | Variable / Indices | Unit | Source (resolution) |
| --- | --- | --- | --- |
| Topographic and climatic | Elevation | Meters | BD ALTI v.2.0: digital terrain model (25 m) |
|  | Slope | Percent | BD ALTI v.2.0: digital terrain model (25 m) |
|  | Cosinus(Aspect) |  | BD ALTI v.2.0: digital terrain model (25 m) |
|  | Sinus(Aspect) |  | BD ALTI v.2.0: digital terrain model (25 m) |
|  | Mean annual temperature | Celsius | Worldclim v2 (1 km) |
|  | Mean annual precipitations | Millimeters | Worldclim v2 (1 km) |
| Forest composition | Proportion of broadleaves | Percent | BD Forêt v2 (0.5 ha) |
|  | Proportion of conifers | Percent | BD Forêt v2 (0.5 ha) |
|  | Proportion of mixed forests | Percent | BD Forêt v2 (0.5 ha) |
| Forest diversity (composition) | Shannon diversity of forest types |  | BD Forêt v2 (0.5 ha) |
|  | Simpson diversity of forest types |  | BD Forêt v2 (0.5 ha) |
| Forest cover | Proportion of forest cover | Percent | BD Forêt v2 (0.5 ha) |
| Forest connectivity (forest patches) | Contagion index | Percent | BD Forêt v2 (0.5 ha) |
|  | Cohesion index | Percent | BD Forêt v2 (0.5 ha) |
|  | Aggregation index | Percent | BD Forêt v2 (0.5 ha) |
| Forest heterogeneity | Shannon diversity of forest patches |  | Corine Land Cover (25 ha) |
|  | Simpson diversity of forest patches |  | Corine Land Cover (25 ha) |
| Forest fragmentation | Landscape shape index |  | OSO (10 m) |
|  | Edge density | Meters / hectare | OSO (10 m) |

### Forest variables

We used QGIS (version 3.16.6-Hannover) and R v 4.5.0 (R Core Team 2025) to extract forest characteristics in the different buffers around presence and background points locations. We extracted forest composition summarized as the proportion of broadleaves, conifers and mixed forests types from the BD Forêt version 2.0 (National Forest Inventory, https://inventaire-forestier.ign.fr/IMG/pdf/dc_bdforet_2-0.pdf, Table 2).

We then calculated Shannon and Simpson diversity indices based on the forest types issued from the BD Forêt v2 (27 forest types in total, see Inventaire Forestier National 2008). Both indices account for the number of types and their abundance within each buffer, they vary from 0 (only 1 type within the buffer) to plus infinite (types equitably distributed within each buffer). The Shannon index puts higher weigh on rare elements while the Simpson index weighs more common elements (Table 2).

We finally calculated the forest cover within each buffer (Table 2).

### Landscape variables

Using the R package landscapemetrics (Hesselbarth et al. 2019), we calculated different landscape variables that account for the different approaches in landscape ecology: the patch characterizing a homogeneous unit, the class characterizing all patches belonging to the same type and the landscape characterizing all patches and classes within a given study window (Sertel et al. 2018). We first calculated forest connectivity indices using the BD Forêt v2 taking into account forest cover only (see McGarigal 2012):

i. the contagion index measures the probability that two patches belong to the same class. The index varies from 0 (a lot of dispersed patches) to 100% (large contiguous patches);
ii. the patch cohesion index measures the connectivity of the patch type studied. The index also varies from 0 (when the studied class is fragmented) to 100% (when the studied class is aggregated);
iii. the aggregation index is calculated using a contiguity matrix and measures, for each combination of patches, the frequency at which patches appear contiguous within a given window. The index varies from 0 (when the landscape is totally fragmented) to 100% (when the landscape is composed of fully contiguous patches, He et al. 2000).

Second, we calculated indices characterizing forest landscape heterogeneity: Shannon and Simpson indices measure the forest landscape diversity accounting for the number of patch classes and their abundance. Both indices were calculated from Corine Land Cover 2018 (https://land.copernicus.eu/pan-european/corine-land-cover/clc2018, Büttner et al. 2021).

Third, we calculated several forest fragmentation indices based on the forest cover layer of CES-OSO (land-use map produced by the National centre of expertise, https://artificialisation.developpement-durable.gouv.fr/bases-donnees/oso-theia):

i. the landscape shape index measures the shape of the study class (forest cover) compared to a maximal level of aggregation within this class. The index varies from 1 (when the landscape is composed of only one forest patch) to the infinite;
ii. the edge index measures edge length per surface unit (meters per hectare of forest patches in our case, calculated for each buffer). The index varies from 0 (no edge) to infinite.

### Biogeoclimatic variables

We extracted biogeoclimatic variables to contextualize the relationships between stand and landscape variables and the occurrence of Black Woodpecker cavities. For each location, we extracted the elevation, slope, aspect (cosinus and sinus transformed) based on the digital terrain model (BDAlti V2.0), mean annual temperature and precipitations issued of Worldclim 2 (Fick and Hijmans 2017, Table 2). For these variables only, we limited extraction to the location of cavity presences and background points, and not to buffers, since we used them to contextualize the relationships between landscape variables and presence of cavities.

### Statistical analyses

We used R v4.5.0 (R Core Team 2025) for all analyses. Following Zuur et al. (2010), we explored the data for variance homogeneity and unaccounted sources of variation, outliers, collinearity and interactions with biogeoclimatic variables. We used linear mixed model to analyze the occurrence of Black Woodpecker cavities as a function of the different explanatory variables listed above (Table 2). We fitted models using the function glmmTMB from the glmmTMB package (Magnusson et al. 2017) with binomial error distribution for occurrence data (presences and background points), logit link and site as random effect to account for the fact that observations from the same site may be more similar than observations issued from two different sites. Before processing for model selection, we used the Variance Inflation Factor (VIF, package car, Fox and Weisberg 2019) to detect multicollinearity amongst explanatory variables and excluded variables with a VIF higher than 3 points, a more selective threshold than the traditional 5 points to exclude collinear explanatory variables. Based on this preliminary selection, we then processed a backward model selection (custom function for model selection in R) based on Akaike Information Criterion (AIC), including all variables with a VIF < 3 points and one-way first-order interactions with biogeoclimatic variables (elevation, aspect, slope, temperature, precipitation). We standardized all variables to compare the magnitudes of the effects. For each scale (10, 100, and 250ha), we discarded variables and interactions until the AIC did not change by more than two points. We tested model hypotheses a posteriori using the package DHARMa (Hartig 2024). All diagnoses revealed good fit of the models (Appendix S1).

## Results

### Model selection on the training dataset

At the 10ha scale, VIF selection kept elevation, slope, aspect (sin and cos), mean precipitation, percentages of conifers and mixed forests, Simpson diversity of forest types, contagion and cohesion index, Simpson diversity of forest patches and landscape shape index. The best model comprised all variables except aspect and interactions (Table 3). All the parameters were significant (p<0.05) except Simpson of forest types. Cohesion, contagion, Simpson of forest patches and percentage of mixed forests had positive effect on cavity excavation, with the highest effect for cohesion (scaled estimate +/-standard error: 1.31 +/- 0.30). Conversely, percentage of conifers, landscape shape index, Simpson of forest types had negative effect with the largest effect for percentage of conifers (−0.29 +/- 0.06). Elevation amplified positive effects of contagion (the higher the elevation, the stronger the contagion effect) but almost canceled the negative effects of Simpson of forest types and landscape shape index (the higher the elevation, the closer to 0 the estimate). Precipitation tended to lessen effects of contagion, percentage of mixed forests and conifers.

**Table 3.**
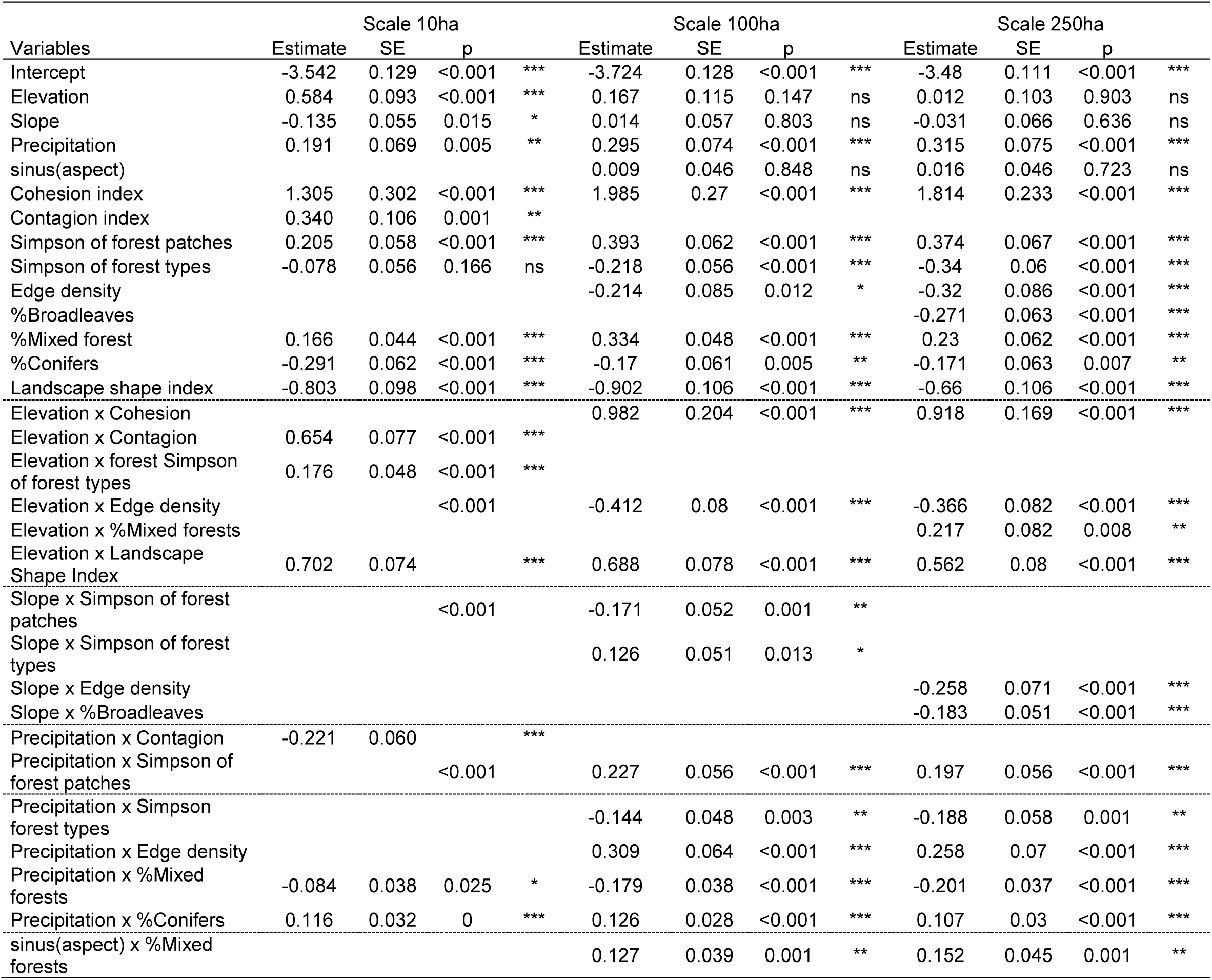
Standardized estimates of the best models predicting the occurrence of Black Woodpecker cavities at three study scales (10ha, 100ha and 250ha). Empty cells correspond to variables not selected in the models. SE: Standard Error. *** p < 0.001; ** p < 0.01; * p < 0.05, (*) p < 0.1.

At the 100ha scale, VIF selection kept elevation, slope, aspect (cos and sin), mean precipitation, percentages of broadleaves, conifers and mixed forests, Simpson of forest types, edge density, cohesion, landscape shape index and Simpson of forest patches. The best model kept all variables except aspect (cosinus), percentage of broadleaves and interactions. All parameters were significant. Cohesion, percentage of mixed forests and Simpson forest patches had positive effects, with higher magnitude for cohesion (1.99 +/- 0.27), but also larger estimates than at the 10ha scale (notably twice as double for Simpson and percentage of mixed forest, Table 3). Percentage of conifers, edge density, Simpson of forest types and landscape shape index all had negative effects on cavity occurrence, with the higher magnitude for landscape shape index (−0.90 +/- 0.11). Elevation amplified the positive effect of cohesion and the negative effects of edge density, but lessened the effect of landscape shape index. Slope lessened both Simpson indices effects. Precipitation amplified both Simpson effects but lessened the negative effects of edge density and conifers and the positive effect of broadleaves. Finally, effects of percentage of mixed forests were higher on south-exposed slopes (high values of sinus aspect)

At the 250ha scale, VIF selection kept the same variables as for the 100ha scale, namely elevation, slope, aspect (cos and sin), mean precipitation, percentages of broadleaves, conifers and mixed forests, Simpson of forest types, edge density, cohesion, landscape shape index and Simpson of forest patches. The best model comprised all these variables except for cosinus(aspect). All parameters were significant except elevation, slope and sinus(aspect). Results were globally comparable to the 100ha scale, with positive effects of cohesion, percentage of mixed forests and Simpson of forest patches, and higher magnitude for cohesion (1.81 +/- 0.23). Conversely, edge density, percentage of broadleaves and conifers, Simpson of forest types and landscape shape index had negative effects with a higher magnitude for landscape shape index (−0.66 +/- 0.11). Elevation amplified the positive effects of cohesion and percentage of mixed forests, and the negative effects of edge density, but lessened the negative effect of landscape shape index. Slope amplified the negative effects of edge density and percentage of broadleaves. As for the 100ha scale, Precipitation amplified both Simpson effects but lessened the negative effects of edge density and conifers and the positive effect of broadleaves. Finally, as for the 100ha scale, effects of percentage of mixed forests were higher on south-exposed slopes (high values of sinus aspect).

Note that in general, we observed higher magnitudes at the 100ha scale, which correspond to the core home range of the Black Woodpecker. Also, interactions had lesser effects on magnitude than univariate effects and some adverse interaction nearly cancelled or reversed the single effects observed (e.g. higher precipitation reduced the negative effect of edge density both at the 100 and 250ha scales, Table 3).

## Discussion

### Mixed broadleaves conifers forests favour cavity excavation by the Black Woodpecker

We partially validated the hypothesis that the presence of both broadleaves and conifers favoured the excavation of cavity by the Black Woodpecker. Indeed, we showed a sharp increase in occurrence probability with increasing percentage of mixed forest (Table 3). It appears that such mixing is more favourable to cavity excavation than juxtaposed pure broadleaves and conifer forests patches, contrary to what was hypothesised. The proximity of broadleaves and conifers within the same forest patches or stands may represent a way to save energy and improve fitness of the species. Broadleaves are preferentially selected for cavity excavation (Basile et al. 2020; Fernandez and Azkona 1996; Karimi et al. 2018; Kosiński and Kempa 2007) while conifers provide food resources, in particular wood- and carpenter ants (Mikusiński 1997) and saproxylic larvae (Brambilla and Saporetti 2014; Rolstad et al. 1998). In central and western Europe, the Black Woodpecker tends to excavate cavities preferentially in beech (*Fagus sylvatica* L.), while in Scandinavia or parts of central Europe where beech is absent, the bird excavates more on conifers like Scots Pine (Pinus sylvestris L., Angelstam and Mikusiński 1994; Pirovano and Zecca 2014; Zawadzka et al. 2016; Zawadzki and Sławski 2023). Tree species with smooth bark, like beech, would be preferred to prevent predation by the Pine Marten since climbing on such trees species seems more difficult (Kosiński and Kempa 2007; Olano 2015; Puverel et al. 2019; Zahner et al. 2017). In the Alps, beech stands would also be favoured because they present larger mean diameter due to a less intensive harvesting and lower densities compared to other forests (De Rosa et al. 2016). The Black Woodpecker may avoid excavating cavities in coniferous stands with low branches, that also favour predation (Bocca et al. 2007). This is confirmed by our results on forest composition at different scales. At all scales, the proportion of conifers had a negative effect on the occurrence of cavities. At the 250ha scale, which corresponds to the extended home range, the effect of broadleaves was negative, and Simpson for composition had increasing negative effect across the scales, indicating decreasing occurrence probability in landscapes with homogeneous forest composition. The Black Woodpecker generally avoids conifer-dominated stands and landscapes, but also pure broadleaves landscapes devoid of conifers at a large scale.

### Black Woodpecker cavities occur more in unfragmented forest landscapes

Cohesion positively affected the presence of Black Woodpecker cavities occurrence, while e.g. edge density or landscape shape index had negative effects, which validated our second hypothesis. Indeed, the occurrence of cavities decreased significantly with increasing edge abundance, and was almost null when the proportion of edge reached 100m per hectare (Table 3). We observed the larger impact of edges at the 100ha scale, which shows that, in our Western-European context, the Black Woodpecker needs large continuous forest areas to excavate cavities in its core home range, as confirmed by the results on forest cover. The strong positive effect of the cohesion index at the three scales confirmed this statement (Table 3). While the Black Woodpecker may cover large distances for foraging, it seems that large unfragmented forest areas are necessary for cavity excavation (Bełcik et al. 2025). However, this result does not seem to be generalizable over the whole distribution of the species. Indeed, in Sweden, Tjernberg et al. (1993) showed that the Black Woodpecker also occurs in fragmented forest landscapes, as long as it can fly from one feeding ground to another. Thus, it can live in landscapes with as low as 26% of forested areas (Dorresteijn et al. 2013) which confirms the contextualised response of the species to fragmentation. Zawadzki and Sławski (2023) also found breeding cavities either within forest stands, green-tree retention patches and isolated trees, but with a higher occurrence within stands.

Similarly, we also validated our third hypothesis on forest heterogeneity (as measured by distribution and size of forest patches). Indeed, we showed that increasing Simpson diversity of forest patches favoured the occurrence of the Black Woodpecker cavities, although the magnitude of this effect was relatively low, compared to previous forest landscape metrics. As shown by the previous results on forest configuration, this confirms that the Black Woodpecker does not need patchy open and forested landscape to excavate in our context, but rather homogeneous forest landscapes with limited edges.

### The probability of cavity excavation is context dependant

Elevation, mean annual precipitation and more marginally slope and aspect were often selected as variables characterizing the general context as well as in the interactions of the model. While occurrence of cavities generally increased with elevation and precipitations across all scales, these variables also had a various effects on the interactions with other variables, notably landscape metrics (Table 3). For example, the effects of cohesion were even higher at high elevation at 100ha and 250ha scales. Integrating context in the relationships between the presence of cavities and forest characteristics may help to better understand favourable excavating conditions, as well as adapt forest management and planning to it. These effects may also reflect other biotic and abiotic factors not included in these analyses, like e.g. mean tree diameter, local tree species composition or stand density, since large trees are necessary for the Black Woodpecker to excavate a suitable cavity (e.g. Basile et al. 2020).

### Limitations and research perspectives

The dataset we analysed was, by essence, heterogeneous since it originated from management and naturalist prospections without a real systematic approach. First, it is unbalanced between the different sites and dominated by the Cevennes dataset. The random site effect we included in the models partially cope for this discrepancy, but it would be worthwhile to complete the inventories to balance more such a dataset. To our knowledge, it is however the larger dataset used for such analyses so far.

Second, it would have ideally been worth verifying a larger sample of the data in the field than the 80 cavities we checked. However, even with this small control sample, we showed that the maps provided were relevant since we found the majority of the cavities mapped in our database. Cavity inventories without pole cameras to check for potential cavity initiations generate false positives in the models that may bias the results and could be taken into account by adding a detection probability term (aka. N-mixture models). Given the total number of observations, verifying all the cavities would have been impossible, and we also assume that the bias remains marginal and not systematic, since the birds very probably initiate cavities in conditions similar as for full cavities excavation. Some observers may have also indirectly controlled the effective presence of cavities by observations of nesting animals or insects, but such validation was not systematically reported. In addition, model validation showed robustness of the model (Appendix S1), but estimations would be more precise with full cavity inventories. Such modelling approach would also be more informative with active nesting and roosting cavities, since it would help documenting the bird’s biology more precisely. But at such a scale, this would require huge fieldwork to have a dataset allowing the analysis. We think our approach is a good compromise regarding the value of the compiled dataset, and remains informative regarding the role of cavities for other species than the Woodpecker itself.

We voluntarily excluded dead trees from the dataset, in order to limit the noise due to other uncontrolled factors like tree dieback. We were conscious that this may have biased the results since dead trees and snags are an important part of woodpecker foraging and breeding niche (Angelstam et al. 2002; Rolstad et al. 1998). It also seems that this habitat is more used in conifer-dominated regions, such as eastern and northern Europe (e.g. Zawadzki and Sławski 2023) where other more favourable tree-species are scarce. In addition, the managed forests we worked in (except a few plots in the Alps) are generally very poor in standing dead trees and snags of acceptable dimensions (e.g. Paillet et al. 2015). It is probable (but difficult to verify) that living trees were more systematically observed than dead trees in the inventory process. For this methodological reason at least, it seems more reasonable to exclude dead trees and snags being conscious that this may bias our results.

More generally, we could not analyse the effects of forest structure and precise composition at different scales around cavities observations, notably the degree of stand maturity and the quantity of deadwood at the large scale. Such data would have been interesting to specify the needs of the species, notably in terms of food resources linked to deadwood. These variables could be derived from airborne LiDAR interpretation (Fuhr et al. 2022; Zellweger et al. 2013), but such data were not available across all the sites we studied, and would have required additional field data to calibrate LiDAR models. In a near future, the ongoing French nation-wide LiDAR acquisition program could help completing the present study, using the same processes of data selection, as well as specifying the requirements of the Black Woodpecker in terms of forest structure. Also, forest composition could be assessed using multispectral data such as Sentinel 2 (e.g. Grabska et al. 2019). More precise mapping of forest and landscape attributes could also be of use in the study of cavity excavation by the Black Woodpecker. For example, mapping issued of the BD Forêts v2 has a resolution of 0.5ha, which may cause noise for the results at the 10ha scale, especially for landscape indices (such as fragmentation, contagion, etc). However, it was the most precise data systematically available for the whole dataset.

Finally, generalisation of our approach in other locations of the distribution range of the species would be interesting, since its ecological preferences may vary with the biogeographical context. In turn, all this would help the preservation of this keystone species for the forest ecosystem, and the cavities excavated that support numerous secondary species from other taxa.

### Conclusions and implications for forest management

We studied the forest composition and landscape variables that influenced Black Woodpecker cavity occurrence. From a strict forest management point of view, landscape variables are rarely under the control of forest management. However, some results may be transferred to management planning at the forest or a larger scale. At the 10ha (management unit) scale, we showed that forest composition (mixed forests) had significant effects, while at larger 100ha (core home range) and 250ha (home range) scales, fragmentation and connectivity had stronger effects. Therefore, maintaining and favouring mixed stands and avoiding pure conifer stands locally would be beneficial to the Black Woodpecker and all the species that depend on its cavities for living. Mixed forests have proved to be beneficial both for biodiversity conservation and a large set of other services (e.g. Liang et al. 2016). Such action should then be integrated at the management planning level. At a larger scale, large continuous forested areas confirm the forest character of the Black Woodpecker, but also the need to maintain connectivity at the landscape scale to favour other forest species. Cavity excavation by the Black Woodpecker is also sensitive to edges, as are many other forest bird species, the conservation of which is then conditioned by large unfragmented areas (Robinson et al. 1995; Villard et al. 1999). These recommendations may ultimately be applied at the larger scale, e.g. for a regional or national parks, when forest biodiversity conservation is at stake.

## Acknowledgements

We are indebted to all forest managers, naturalists and other Woodpecker lovers who contributed to cavity mapping. Without their huge work and dedication, this study would not have been possible. B. Algoët, H. Caroff, D. Joud, A. Savine, A. Porte, L. Belenger and B. Guérin especially facilitated the data exchange and interpretation. CC did all the analyses during the Master’s degree training period. We thank the two reviewers of this paper for their constructive comments.

## Declarations Ethical Approval

Not applicable

## Competing interests

We, the authors, declare no competing interests.

## Authors’ contributions

C.C. homogenized the database and calculated all the explanatory variables. C.C. and Y.P. analysed the data and wrote the manuscript text. All authors reviewed the manuscript.

## Funding

Not applicable

## Availability of data and materials

Data and code are available at the following repository: https://entrepot.recherche.data.gouv.fr/dataset.xhtml?persistentId=doi:10.57745/MFIJDB

## Appendix S1: variables correlation and model validation diagnoses using DHARMa

### Variables names

“XX” refers to the scale in ha (10, 100, 250)

*Bioclimatic*: ALTITUDE = Elevation (m); PENTE = slope (deg); cos.ASPECT, sin.ASPECT = cosinus and sinus transformed aspect; TEMP_MEAN = mean temperature (deg); PREC_MEAN = mean precipitation (mm);

*Forest Composition*: FOR_FEUILLU_XX = proportion of broadleaved forest; FOR_RES_XX = proportion of coniferous forest; FOR_MIXTE_XX = proportion of mixed broadleaves / conifers forest; H_DIV_FOR_XX = Shannon index for forest composition; D_DIV_FOR_XX = Simpson index for forest composition

*Landscape*: PLANDXX = proportion of forest; CONNECTXX = patch contagion index; AIXX= patch aggregation index; EDXX = edge density; COHESIONXX = % of patch cohesion; H_DIV_XX = Shannon index for forest patches; D_DIV_XX = Simpson index for forest patches; LSI_XX = landscape Shape Index; EDXX = Edge density.

**Scale 10ha**

**1. Correlations between explanatory variables in the full model**

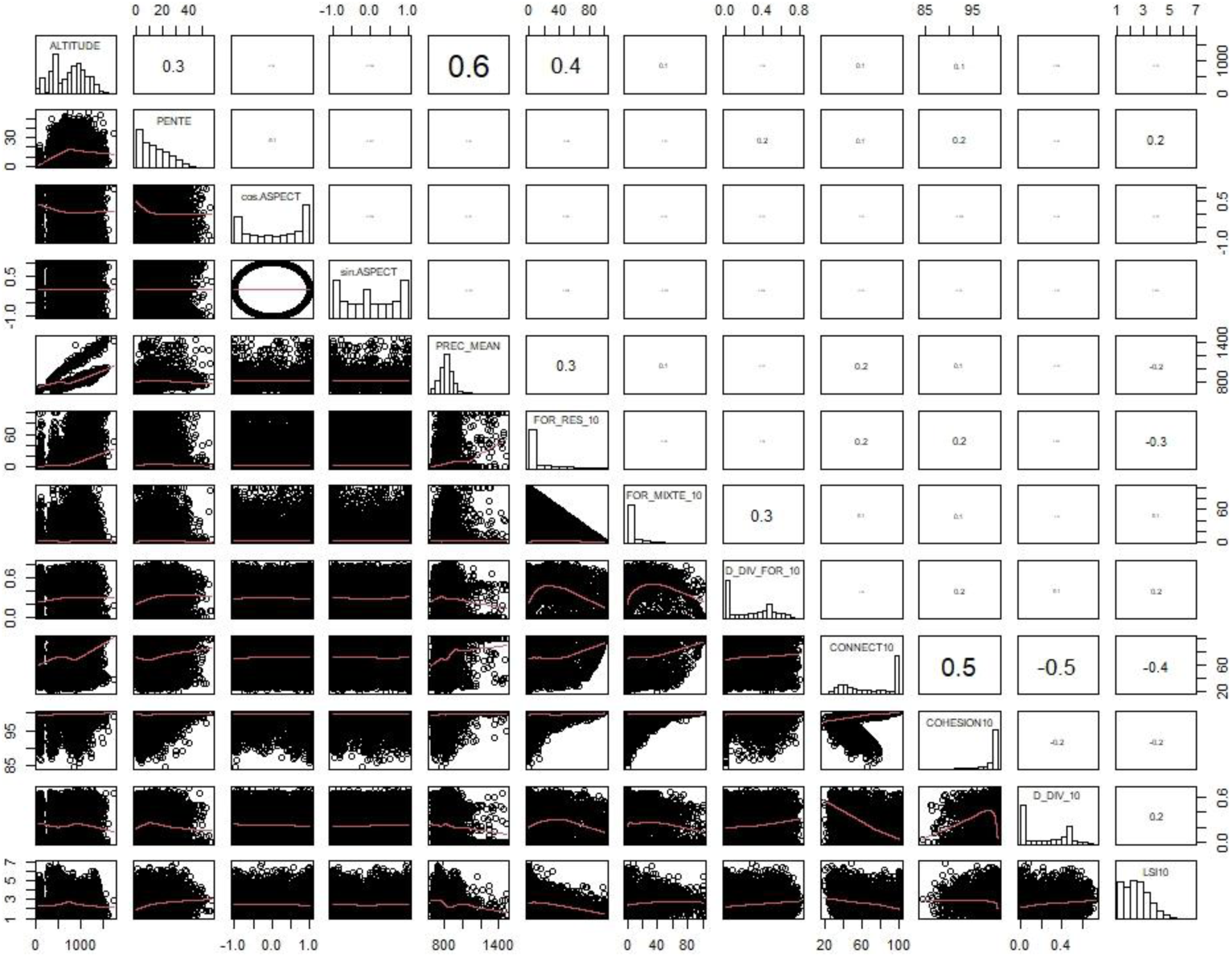

**2. Diagnoses for the model fit**

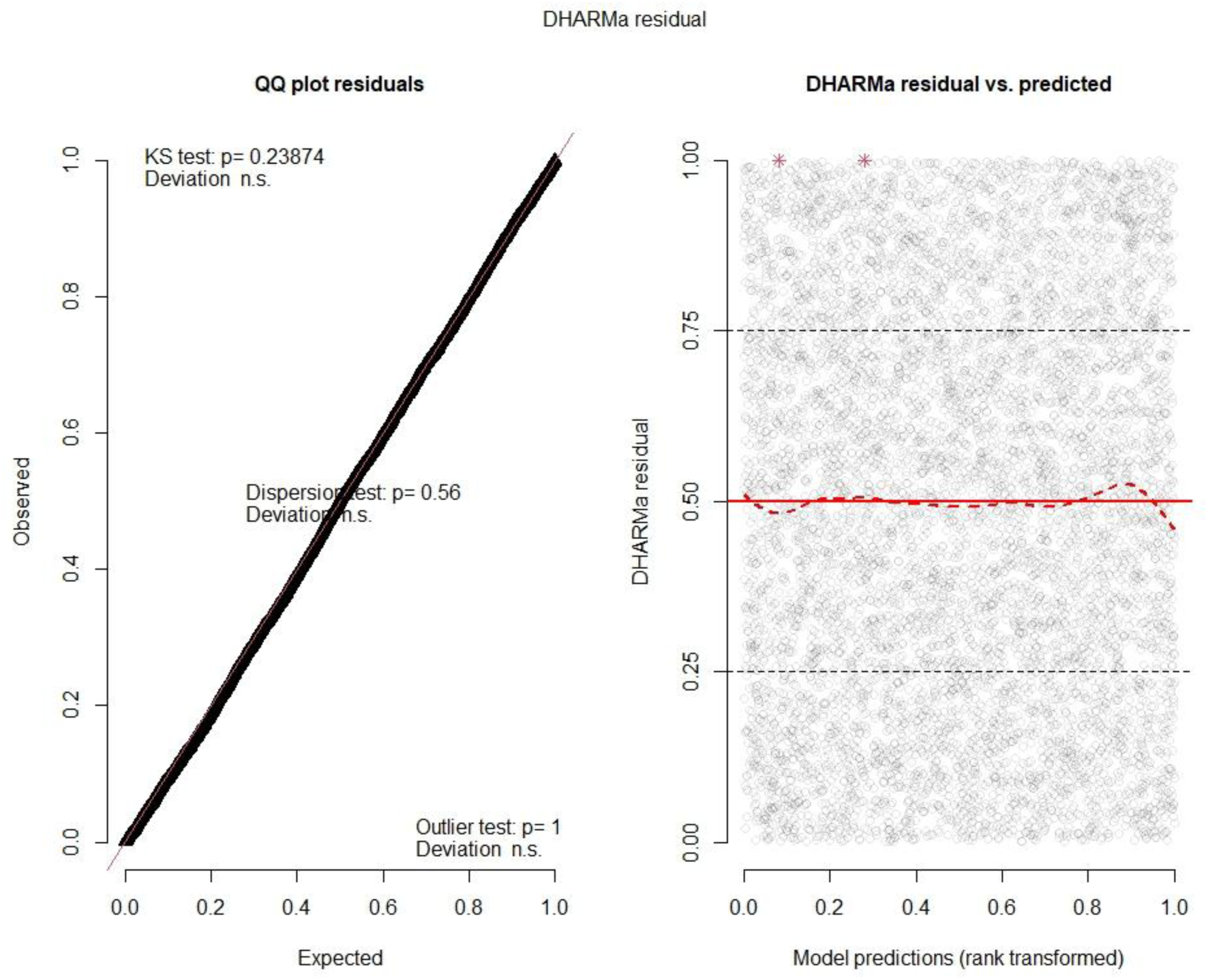

**3. Covariables influence on model fit**

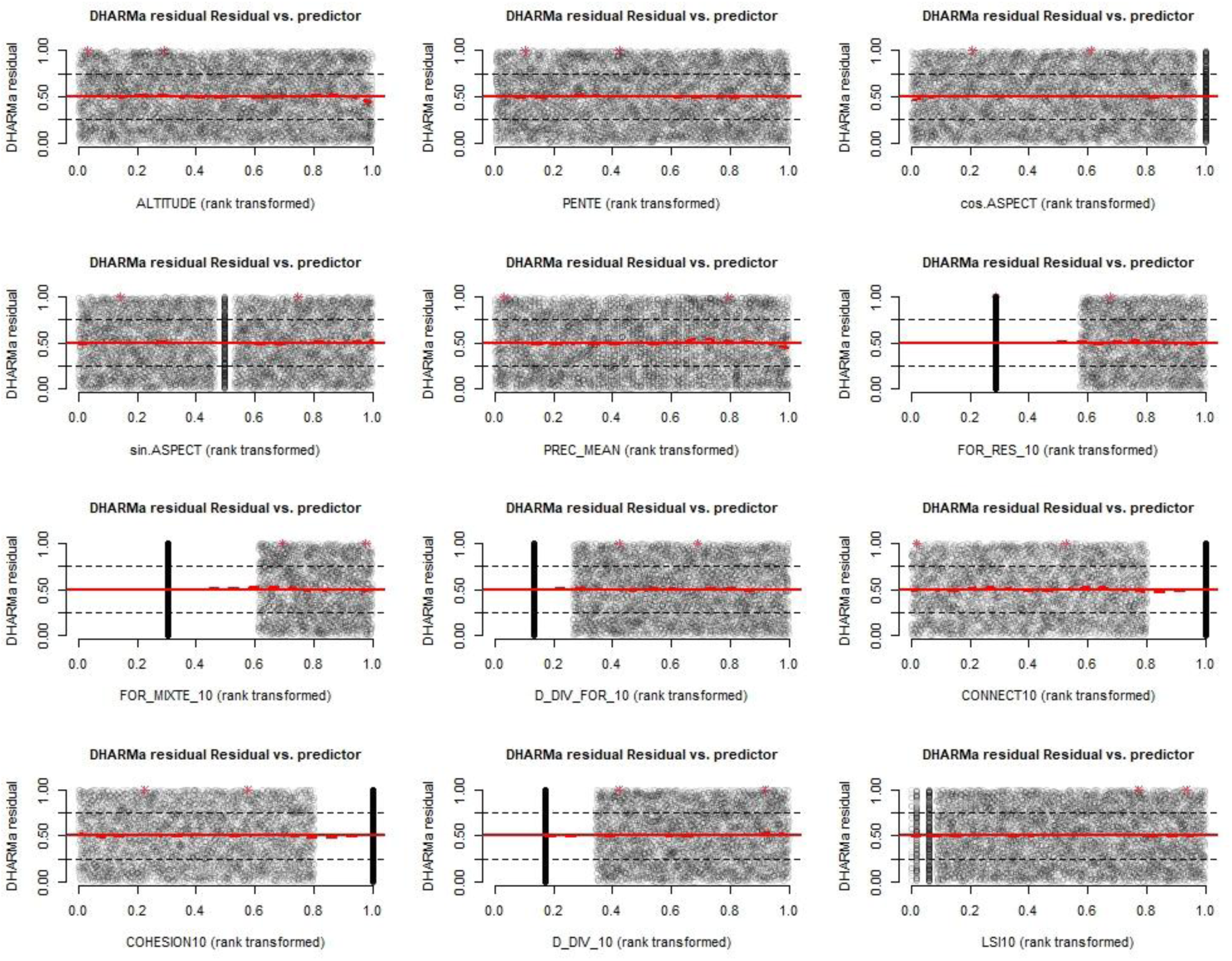

**Scale 100ha**

**1. Correlations between explanatory variables in the full model**

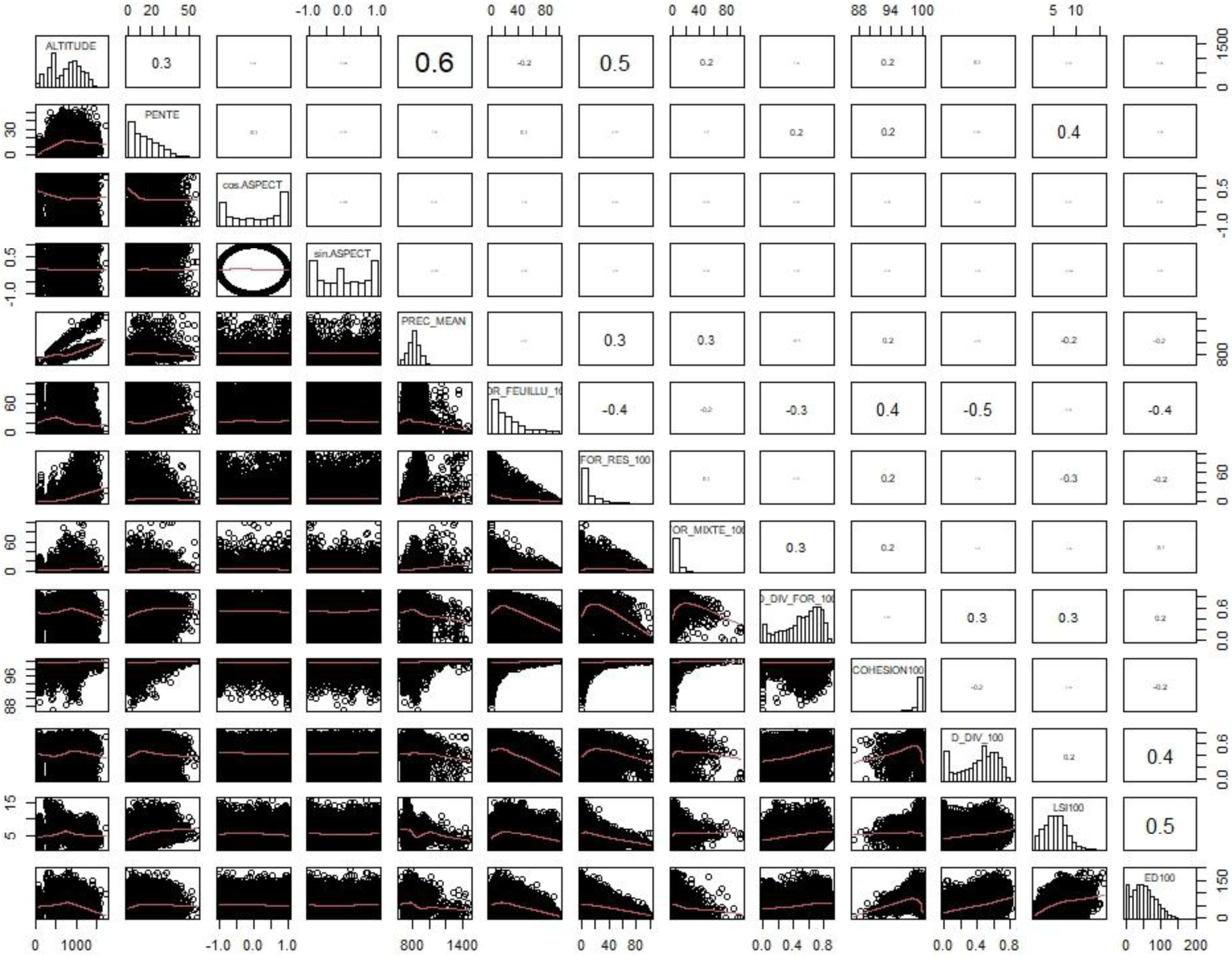

**2. Diagnoses for the model fit**

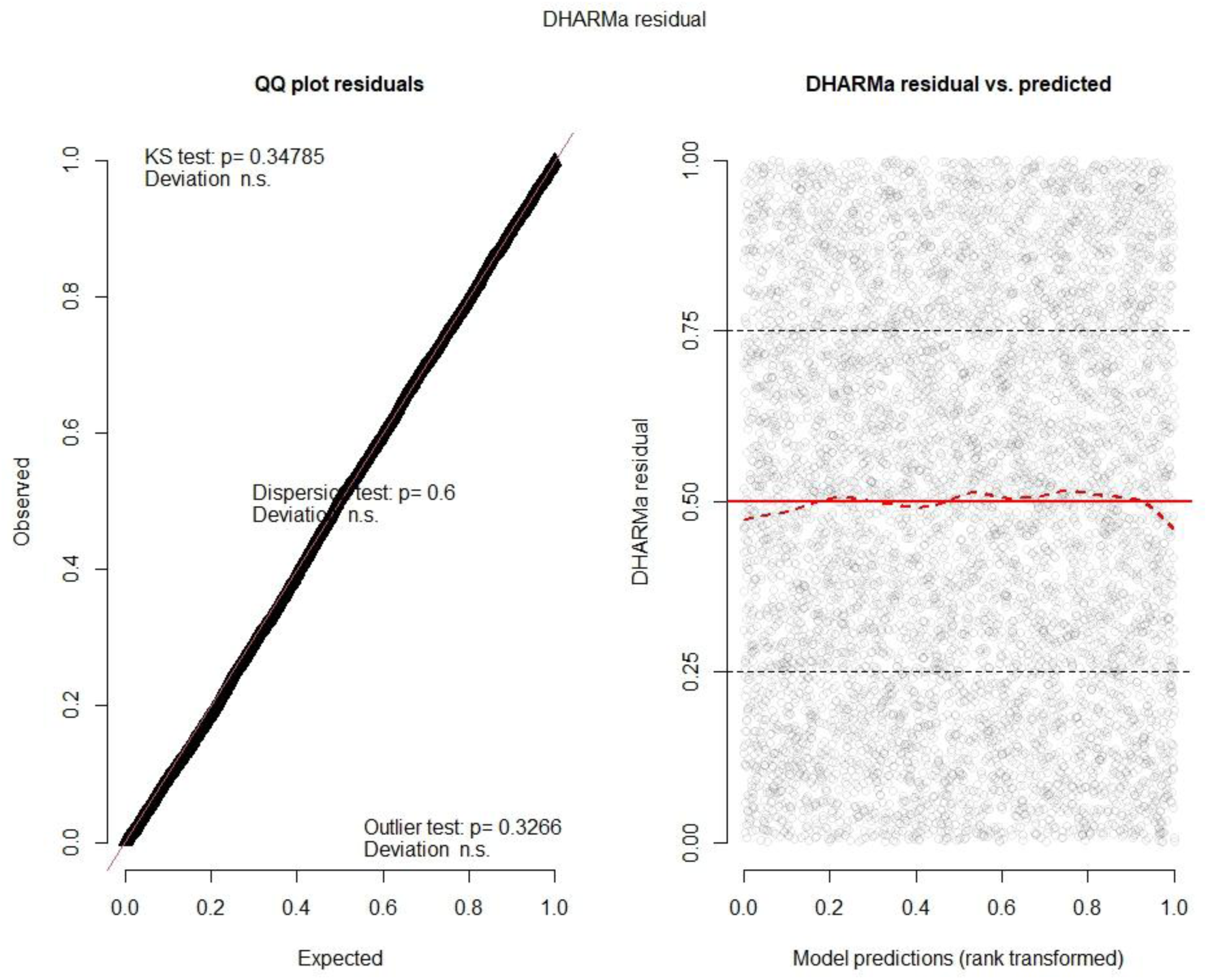

**3. Covariables influence on model fit**

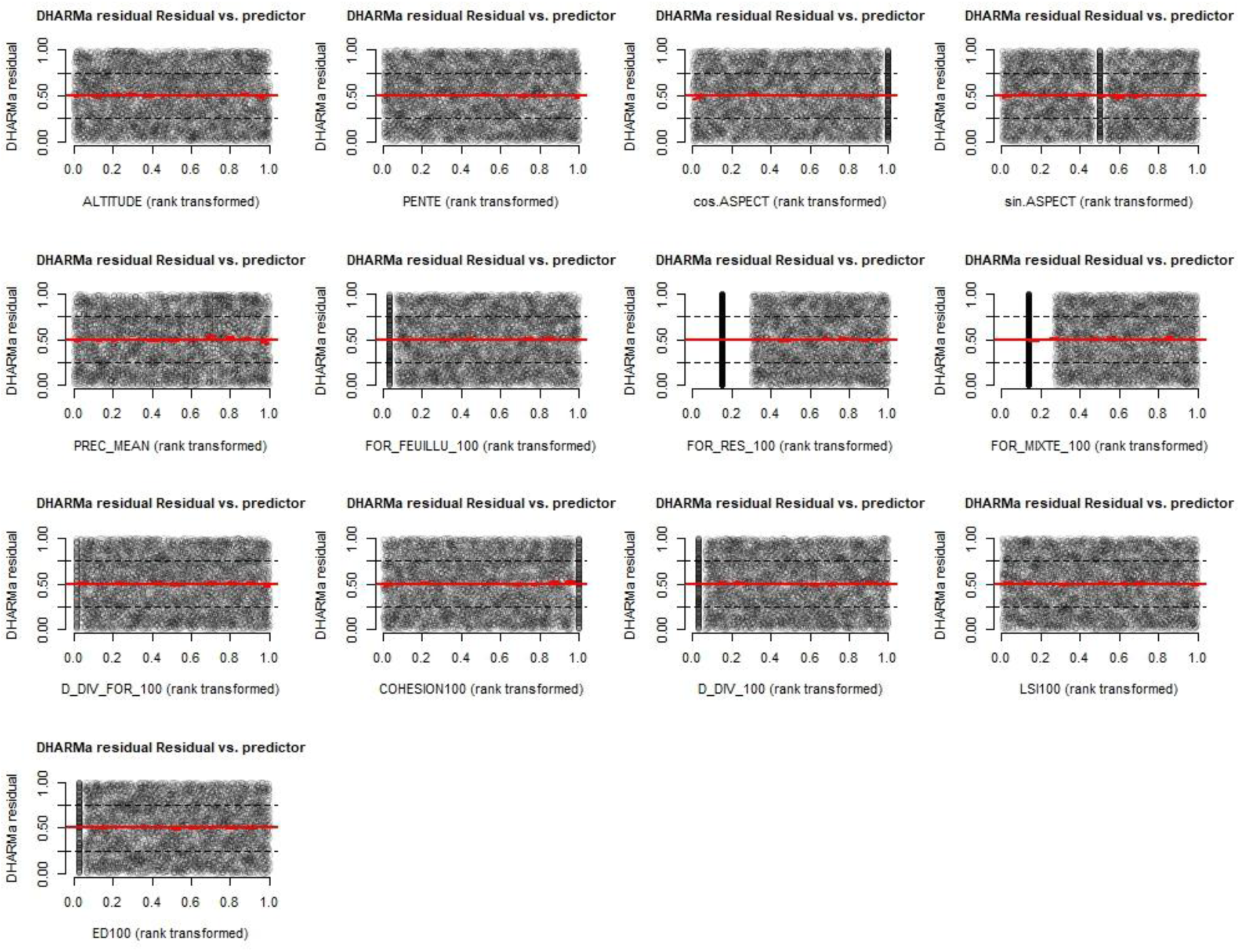

**Scale 250ha**

**1. Correlations between explanatory variables in the full model**

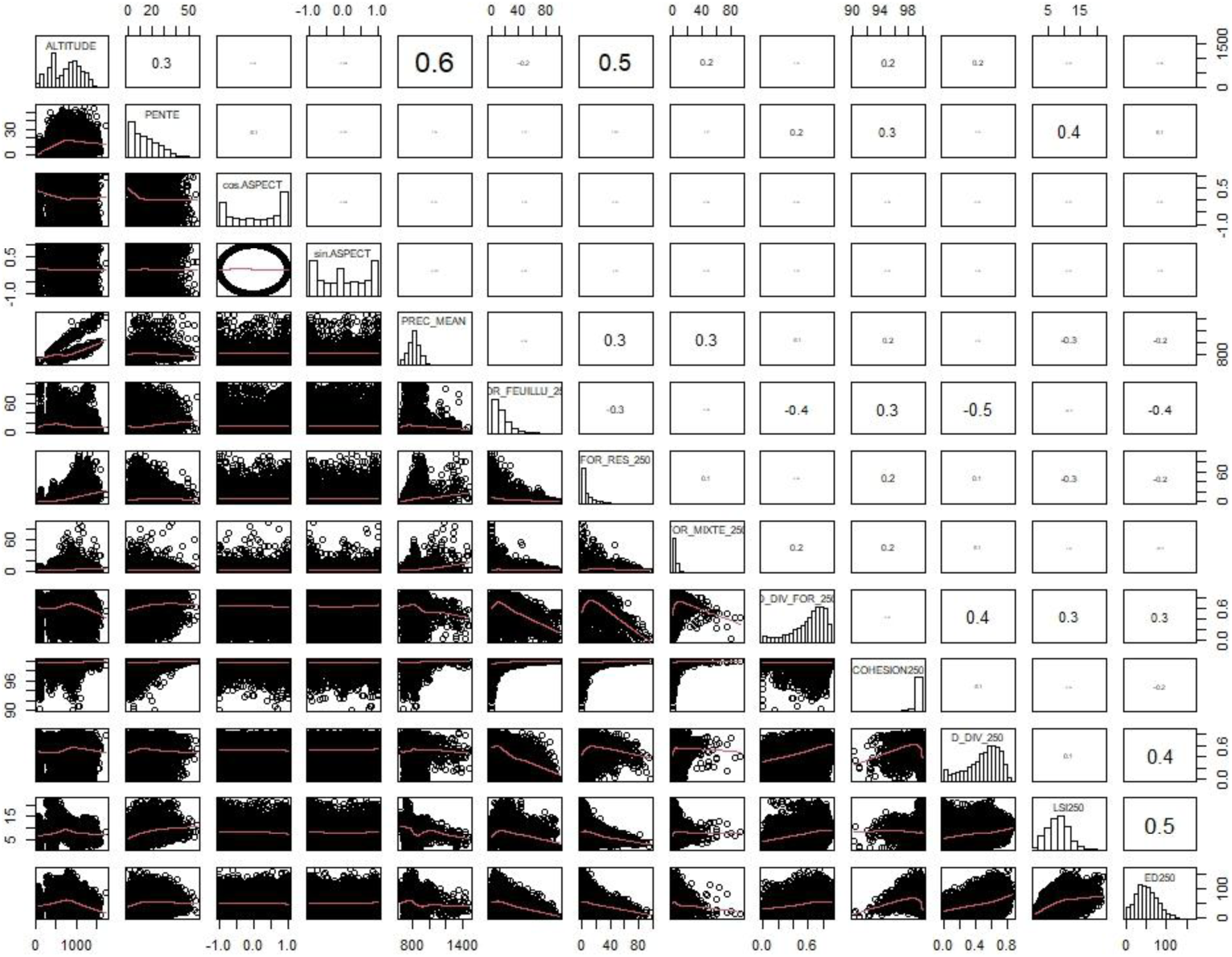

**2. Diagnoses for the model fit**

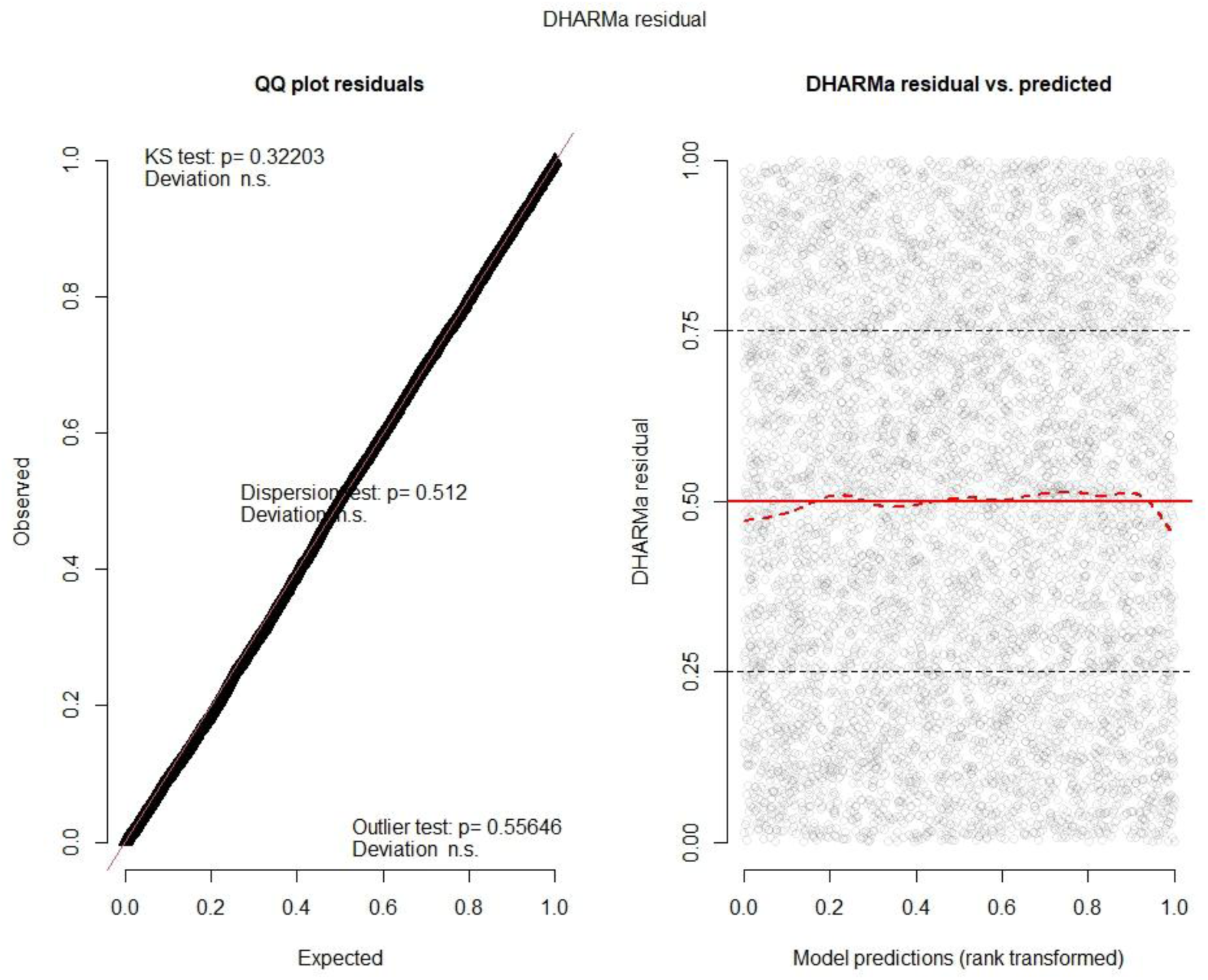

**3. Covariables influence on model fit**

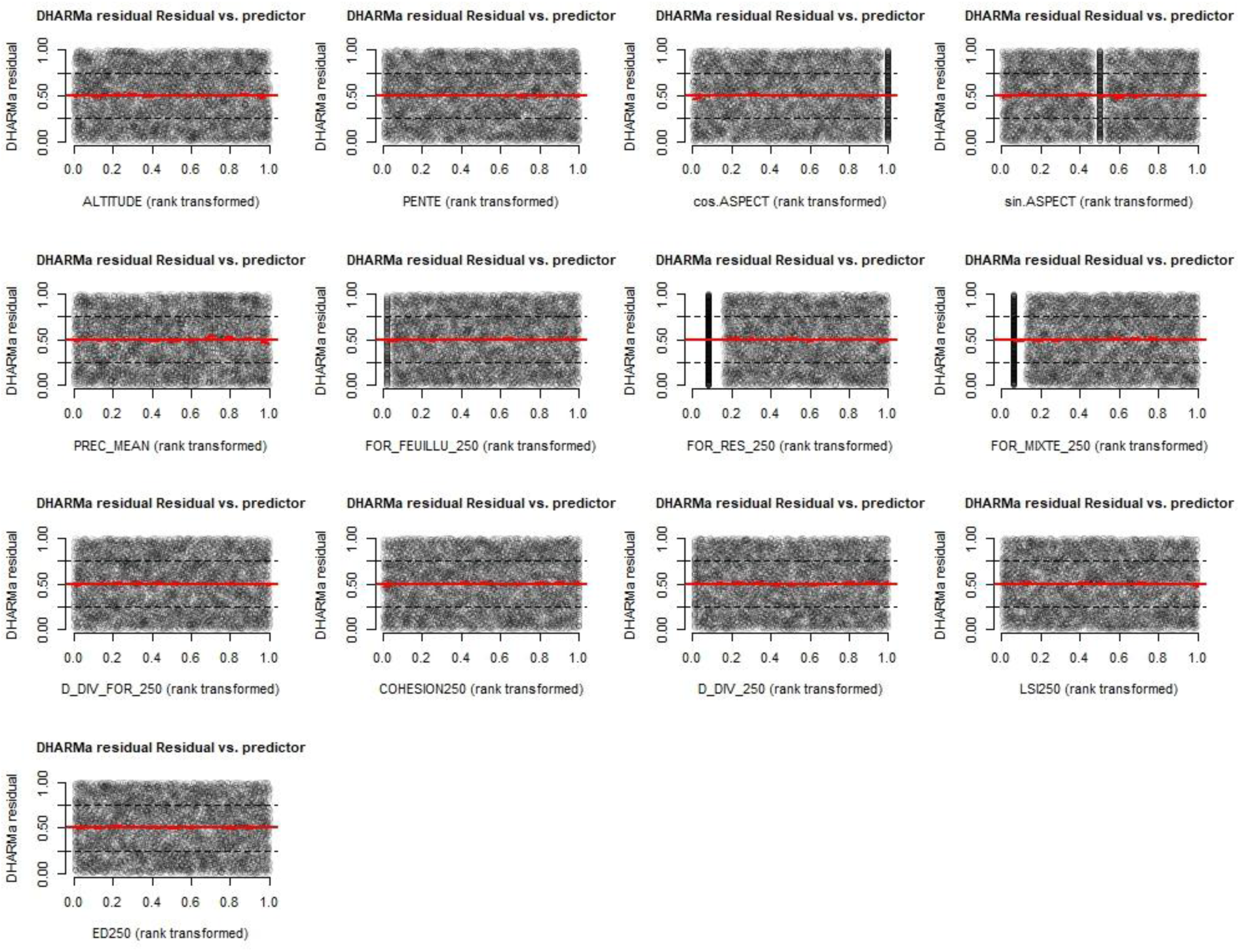

## Notes

https://entrepot.recherche.data.gouv.fr/dataset.xhtml?persistentId=doi:10.57745/MFIJDB

### Competing Interest Statement

The authors have declared no competing interest.

### Summary of Updates

This is the version peer-reviewed and recommended by PCI Ecology. The badge contains the link towards the recommendation.

